# *Reysenbachia aerophila* gen. nov., sp. nov., a facultatively anaerobic, hydrogen-oxidizing, thermophilic bacterium isolated from Kuirau Park, Rotorua, New Zealand

**DOI:** 10.64898/2026.06.14.732183

**Authors:** M. E. A. Marshall, M. B. Stott, H. E. Welford, K. Lagutin, K. A. Mitchell, C. R. Carere

## Abstract

A facultatively anaerobic, hydrogen-oxidizing, thermophilic bacterium (strain KUI-RB^T^) was isolated from a geothermal spring biofilm in Rotorua, New Zealand. Strain KUI-RB^T^ is a motile, straight rod, measuring approximately 0.7 µm by 1.0 to 1.5 µm with a diderm cell wall. Growth of KUI-RB^T^ occurred from 39 to 74 °C (T_opt_ 64.5 °C), pH 5.0 to 7.5 (pH_opt_ 6.5), and 0 to 1% (w/v) NaCl (NaCl_opt_ 0.4-0.7%, w/v). KUI-RB^T^ utilizes carbon dioxide and various organic carbon substrates as carbon sources and hydrogen as an electron donor. KUI-RB^T^ can use oxygen (0-21%, v/v), elemental sulfur, thiosulfate, sulfite, nitrate, arsenate, and selenate as terminal electron acceptors. Major fatty acids of strain KUI-RB^T^ include C_20:1_, C_18:1_, and C_18:0_ and the primary quinone is MTK-7. The whole genome G+C content is 34.23 mol%. Phylogenetic analyses indicate KUI-RB^T^ to be a member of the family *Hydrogenothermaceae*, with *Sulfurihydrogenibium azorense* Az-Fu1^T^ its closest characterised relative (94.51% 16S rRNA gene sequence similarity, 78.01% whole genome ANI, 61.34% whole genome AAI). Based on phylogenetic and phenotypic analyses, we propose KUI-RB^T^ represents a novel genus and species within the family *Hydrogenothermaceae*, for which we propose the name *Reysenbachia aerophila* gen. nov., sp. nov. The type strain is KUI-RB^T^ (=KCTC accession =JCM accession).

The GenBank accession number for the 16S rRNA gene sequence of strain KUI-RB^T^ is PZ052650. The GenBank accession number for the whole genome of strain KUI-RB^T^ is JBVODP000000000.

## Introduction

The order *Aquificales* (phylum *Aquificota*), consists of two families, *Aquificaceae* and *Hydrogenothermaceae* (Gupta, 2014; Oren & Garrity, 2021). Characterised representatives within *Hydrogenothermaceae* are thermophilic, neutrophilic, microaerophilic, Gram-negative motile rods capable of sulfur oxidation and carbon dioxide (CO_2_) fixation (Eder & Huber, 2002; Reysenbach *et al*., 2001, 2009). Some, but not all members of the *Hydrogenothermaceae* family perform the ‘knallgas’ reaction, in which hydrogen is coupled with oxygen to form water. Some representatives also grow anaerobically using elemental sulfur, nitrate, arsenate, selenate, or selenite as terminal electron acceptors (Aguiar *et al*., 2004; François *et al*., 2021; Götz *et al*., 2002; Nakagawa *et al*., 2003; Takai *et al*., 2003).

There are currently four characterised genera of *Hydrogenothermaceae*; *Hydrogenothermus* and *Persephonella*, which are primarily detected in marine hydrothermal environments (François *et al*., 2021; Götz *et al*., 2002; Nakagawa *et al*., 2003; Sislak, 2013; Stohr *et al*., 2001), and *Sulfurihydrogenibium* and *Venenivibrio*, which appear to exclusively populate terrestrial geothermal springs (Aguiar *et al*., 2004; Flores *et al*., 2008; Hetzer *et al*., 2008; Nakagawa *et al*., 2005; O’Neill *et al*., 2008; Takai *et al*., 2003). There are five species of *Sulfurihydrogenibium*: *Sulfurihydrogenibium azorense*, *Sulfurihydrogenibium kristjanssonii*, *Sulfurihydrogenibium rodmanii*, *Sulfurihydrogenibium subterraneum* and *Sulfurihydrogenibium yellowstonense* (Table 1) (Aguiar *et al*., 2004; Flores *et al*., 2008; Nakagawa *et al*., 2005; O’Neill *et al*., 2008; Takai *et al*., 2003). *Sulfurihydrogenibium* are frequently detected in terrestrial geothermal springs worldwide, including those in Yellowstone National Park (USA), Ourense (Spain), Azores (Portugal), Nagano Prefecture (Japan), Kunashir Island and Kamchatka Peninsula (Russia) (Dong *et al*., 2019; Escuder-Rodríguez *et al*., 2022; Hamamura *et al*., 2013; Kubo *et al*., 2011; Malygina *et al*., 2023; McKay *et al*., 2022; Merkel *et al*., 2017). In contrast, *Venenivibrio* appears to be restricted to geothermal springs in New Zealand, with no reports from other regions to date (Power *et al*., 2024). There is currently one characterised species of *Venenivibrio* and two candidate species: *Venenivibrio stagnispumantis*, *Venenivibrio* sp. KUI1, and *Venenivibrio* sp. OKO1 (Hetzer *et al*., 2008; Welford *et al*., unpublished).

**Table 1.** Differential characteristics of strain KUI-RB^T^ and related type strains of the family *Hydrogenothermaceae*. Species: 1, *Reysenbachia aerophila* KUI-RB^T^ (this study); 2, *Venenivibrio stagnispumantis* CP.B2^T^ (Hetzer *et al*., 2008); 3, *Sulfurihydrogenibium azorense* AZ-Fu1^T^(Aguiar *et al*., 2004; Nakagawa *et al*., 2005); 4, *Sulfurihydrogenibium kristjanssonii* I6628^T^ (Flores *et al*., 2008); 5, *Sulfurihydrogenibium rodmanii* UZ3-5^T^(O’Neill *et al*., 2008); 6, *Sulfurihydrogenibium subterraneum* HGMK1^T^ (Aguiar *et al*., 2004; Nakagawa *et al*., 2005; Takai *et al*., 2002); 7, *Sulfurihydrogenibium yellowstonense* SS-5^T^ (Nakagawa *et al*., 2005). ND, Not determined.

| Characteristic | 1 | 2 | 3 | 4 | 5 | 6 | 7 |
| --- | --- | --- | --- | --- | --- | --- | --- |
| Temperature range (optimum) (°C) | 39-74 (64.5) | 45-75 (70) | 50-73 (68) | 40-73 (68) | 55-80 (75) | 40-70 (65) | 55-78 (70) |
| pH range (optimum) | 5.0-7.5 (6.5) | 4.8-5.8 (5.4) | 5.5-7.0 (6) | 5.3-7.8 (6.6) | 5.0-7.2 (6.0-6.3) | 6.4-8.8 (7.5) | 6.0-8.0 (7.5) |
| NaCl range (optimum) (% w/v) | 0.0-1.0 (0.4-0.7) | 0.0-0.8 (0.4) | 0.0-0.25 (0.1) | 0.0-0.5 (0.0) | 0.0-0.9 (0.0) | 0-4.8 (0.5) | 0.0-0.6 (0.0) |
| O<sub>2</sub> range (optimum) (% v/v) | 0-21 (21) | 1.0-10.0 (4-8) | 0.2-9.0 (1.8-2.0) | <25 (4-9) | 0.5-14 (4-6) | 0-5 with H<sub>2</sub>, 0-20 with S<sup>0</sup> | 0.1-18 (6) |
| Electron donor | H<sub>2</sub> | H<sub>2</sub> | H<sub>2</sub>, S<sup>0</sup>, SO<sub>3</sub><sup>2-</sup>, S<sub>2</sub>O<sub>3</sub><sup>2-</sup>, Fe<sup>2+</sup> | H<sub>2</sub>, S<sup>0</sup>, S<sub>2</sub>O<sub>3</sub><sup>2-</sup> | S<sup>0</sup> or S<sub>2</sub>O<sub>3</sub><sup>2-</sup> | H<sub>2</sub>, S<sup>0</sup>, S<sub>2</sub>O<sub>3</sub><sup>2-</sup>, Fe<sup>2+</sup> | S<sup>0</sup> or S<sub>2</sub>O<sub>3</sub><sup>2-</sup> |
| Electron acceptor | O<sub>2</sub>, S<sup>0</sup>, S<sub>2</sub>O<sub>3</sub><sup>2-</sup>, SO<sub>3</sub><sup>2-</sup>, NO<sub>3</sub><sup>-</sup>, AsO<sub>4</sub><sup>3-</sup>, SeO<sub>4</sub><sup>2-</sup> | O<sub>2</sub> | O<sub>2</sub>, S<sup>0</sup>, AsO<sub>4</sub><sup>3-</sup>, Fe<sup>3+</sup> | O<sub>2</sub> | O<sub>2</sub> | O<sub>2</sub>, NO<sub>3</sub><sup>-</sup>, Fe<sup>3+</sup>, AsO<sub>4</sub><sup>3-</sup>, SeO<sub>4</sub><sup>2-</sup>, SeO<sub>3</sub><sup>2-</sup> | O<sub>2</sub> |
| Carbon source | CO<sub>2</sub>, Bacto peptone, trypticase peptone, Casamino acids, sucrose, glucose, starch, formate, citrate, succinate, acetate, pyruvate | CO<sub>2</sub> | CO<sub>2</sub>, yeast extract, Bacto peptone, trypticase peptone, Casamino acids | CO<sub>2</sub>, succinate | CO<sub>2</sub> | CO<sub>2</sub>, acetate | CO<sub>2</sub>, yeast extract, Bacto peptone, trypticase peptone, Casamino acids, sucrose, glucose, starch, formate, citrate, acetate, propionate |
| DNA G+C content (mol%) | 34.2 | 29.2 | 33.6 | 28.1 | 35 | 31.3 | 32 |
| Genome size (Mb) | 1.62 | 1.63 | 1.64 | ND | ND | 1.61 | 1.53 |

Here we report the phylogenetic, phylogenomic, and physiological characterisation of a novel strain, strain KUI-RB^T^, isolated from a terrestrial geothermal spring in Rotorua, New Zealand. Based on our findings, we propose that strain KUI-RB^T^ represents a novel species of a new genus within the family *Hydrogenothermaceae,* for which the name *Reysenbachia aerophila* gen. nov., sp. nov. is proposed.

## Methods and Results

### Sample collection, enrichment, and purification

A biofilm sample was collected in April 2024 from overflow between two geothermal features within Kuirau Park, Rotorua, New Zealand (-38.129684, 176.243673). The temperature of the sampling location was 55 °C and the pH was 8.2. Approximately 10 mL of biofilm and spring water was collected in a 50 mL falcon tube, transported at ambient temperature, then stored at 4 °C for three days before processing.

The biofilm sample was vortexed, and 1 mL of the resulting suspension was inoculated into a 60 mL serum bottle containing 25 mL of modified MJTSO medium (JCM 446) (pH 7.8). MJTSO medium contained (per litre): 3 g NaCl, 0.34 g MgCl_2_·6H_2_O, 0.42 g MgSO_4_·7H_2_O, 0.05 g KCl, 0.25 g NH_4_Cl, 0.07 g CaCl_2_·2H_2_O, 1 g Na_2_S_2_O_3_·5H_2_O, 0.1 g yeast extract, 3 g elemental sulfur (S^0^), 0.14 g K_2_HPO_4_, 1 g NaHCO_3_, and 10 mL of trace mineral solution. Trace mineral solution contained (per litre): 500 mg Na-EDTA·2H_2_O, 150 mg CoCl_2_·6H_2_O, 100 mg MnCl_2_·4H_2_O, 100 mg FeSO_4_·7H_2_O, 100 mg ZnCl_2_, 40 mg AlCl_3_·6H_2_O, 30 mg Na_2_(WO)_4_·2H_2_O, 20 mg CuCl_2_·2H_2_O, 20 mg NiSO_4_·6H_2_O, 10 mg NaSeO_3_, 10 mg H_3_BO_3_, and 10 mg Na_2_MoO_4_·2H_2_O. All medium components, except S^0^, K_2_HPO_4_, and NaHCO_3_ were combined, then autoclaved at 121 °C for 20 min at 15 psi. Stock solutions of filter-sterilized (0.22 µm) K_2_HPO_4_ and NaHCO_3_ were aseptically added after autoclaving, and the medium was adjusted to pH 7.8 with 2.5% (v/v) HCl. S^0^ was autoclaved within individual serum bottles at 105 °C for 18 min at 15 psi. Medium was then added to these S^0^-containing serum bottles, which were sealed with butyl rubber stoppers and flushed with 100% (v/v) N_2_ for 12 seconds/mL of medium. Prior to inoculation, the headspace composition was adjusted to 76% H_2_, 19% CO_2_, 5% O_2_ (v/v).

Enrichment cultures were subcultured twice (10%, v/v) following an initial six-day incubation period. Subsequently, three consecutive rounds of serial dilution to extinction were performed, resulting in the isolation of strain KUI-RB^T^. All incubations were conducted at 55 °C without shaking.

Unless stated otherwise, all characterisation experiments were conducted with agitation at 55 °C using 10 mL of MJTSO medium (pH 6 to pH 7) and a 17 mL headspace comprising 76% H_2_, 19% CO_2_, and 5% O_2_ (v/v). Elemental sulfur was omitted from the medium during characterisation, as growth remained consistent in its absence and this eliminated spectrophotometric interference in optical density measurements. Growth was assessed in triplicate by measuring optical density at 610 nm (OD_610_) using a DR900 colorimeter (Hach, Loveland, USA) and, when necessary, by direct cell counts using a Helber counting chamber (Hawksley, Brighton & Hove, UK). All phenotypic observations were confirmed via two sequential subcultures at the prescribed condition. Culture purity was routinely confirmed by 16S rRNA gene sequencing using 2× Myfi Mix (Meridian Bioscience, Cincinnati, USA) and primers 9F (5’-AGAGTTTGATCMTGG-3’) and 1492R (5’-GGHTACCTTGTTACGA-3’) (0.4 µM) (Weisburg *et al*., 1991).

### Growth characteristics

The effect of temperature on the growth of strain KUI-RB^T^ was assessed over the range 36 to 76 °C and growth was observed between 39 and 74 °C (Table 1). To determine the optimal growth temperature and doubling time, OD_610_ measurements were recorded every 4 h across this temperature range using cultures with a headspace of 63% H_2_, 16% CO_2_, and 21% O_2_ (v/v). The doubling time of strain KUI-RB^T^ was 63 min at 66 °C. Maximum specific growth rates were calculated for each temperature and fitted to the Rosso cardinal temperature model with inflection (Fig. S1), extended Ratkowsky model, and Schoolfield model (Ratkowsky *et al*., 1982; Rosso *et al*., 1993; Schoolfield *et al*., 1981). These models converged on an optimal growth temperature (T_opt_) of 64.5 °C (µ_max_: 0.63 h^-1^).

The impact of pH on growth of KUI-RB^T^ was determined via OD_610_ of the culture using MJTSO medium with either a 10 mM tri-sodium citrate/citric acid monohydrate buffer (pH 4.0 to 6.5) or a 12 mM NaHCO_3_ buffer (pH 6.5 to 9.0). The MJTSO medium pH was adjusted after autoclaving by the addition of 2.5% (v/v) HCl or 1 M KOH as appropriate and confirmed with a UP-10 pH/mV Meter (Denver Instrument, Bohemia, USA). Medium pH was similarly measured at the conclusion of the experiment to determine any change in pH due to culture growth. Growth of strain KUI-RB^T^ was observed between pH 5.0 and 7.5 (pH_opt_ 6.5, Table 1). No growth was observed following incubation at pH > 7.5; however, white precipitates were observed in the MJTSO medium and increased in density from pH 7.5 to 9.0. In later organic carbon utilization experiments we noted that strain KUI-RB^T^ grew at pH values from pH 8.0 to 9.0 and without precipitation of MJTSO medium. We therefore conclude that during hydrogenotrophic growth, growth of KUI-RB^T^ is limited to < pH 7.5; however, the range is extended to pH 9.0 under organotrophic conditions. The lack of hydrogenotrophic growth at > pH 7.5 may be due to the loss of an essential nutrient via precipitation with KOH. NaCl tolerance of strain KUI-RB^T^ was assessed between 0 and 2% (w/v). Growth occurred up to 1% (w/v) NaCl, with optimal growth between 0.4 and 0.7% (w/v) (Table 1).

Strain KUI-RB^T^ grew autotrophically using CO_2_ (19% v/v) during initial characterisation. The utilization of organic carbon substrates as carbon sources was assessed in MJTSO medium lacking both yeast extract and S^0^. NaHCO_3_ was used as a buffer for all carbon experiments after control experiments determined strain KUI-RB^T^ did not grow on NaHCO_3_ as a sole carbon source. Organic substrates (0.1%, w/v) were added from sterile stock solutions (yeast extract, Bacto peptone, trypticase peptone, Casamino acids, sucrose, glucose, soluble starch, Na-formate, Na_3_-citrate·2H_2_O, Na_2_-succinate·6H_2_O, Na-acetate, Na-pyruvate and Na-propionate). NaHCO_3_ (1 g L^−1^) was added after autoclaving and the pH was adjusted to 7.8 with 2.5% (v/v) HCl. The headspace consisted of 79% H_2_ and 21% O_2_ (v/v). With the exception of yeast extract and Na-propionate, strain KUI-RB^T^ was able to utilize all substrates tested as carbon sources in place of CO_2_ (Table 1). The pH of the medium following heterotrophic growth ranged from pH 7.8 to 9.0. We also assessed whether KUI-RB^T^ could grow organotrophically in MJTSO medium (pH 7.2) containing trypticase peptone, glucose, Na-formate or Na-acetate (0.1%, w/v) with a headspace of 79% N_2_ and 21% O_2_ (v/v). Strain KUI-RB^T^ was unable to utilize organic carbon in the absence of H_2_.

The use of inorganic electron donors by strain KUI-RB^T^ was assessed in MJTSO medium lacking thiosulfate (Na_2_S_2_O_3_·5H_2_O) and S^0^, and without H_2_ in the headspace (headspace: 79% CO_2_, 21% O_2_, v/v). Potential electron donors (0.1%, w/v) supplied from sterile stock solutions included S^0^, Na_2_S_2_O_3_·5H_2_O, sulfite (Na_2_SO_3_), ammonium (NH_4_Cl), arsenite (NaAsO_2_), or selenite (Na_2_SeO_3_). H_2_ was tested as an electron donor (76% v/v) during initial isolation. Growth was only observed with H_2_ as an electron donor (Table 1).

The utilization of alternative electron acceptors was assessed in MJTSO medium lacking Na_2_S_2_O_3_·5H_2_O and S^0^, and without O_2_ in the headspace (headspace: 80% H_2_, 20% CO_2_, v/v). Potential electron acceptors (0.1%, w/v) supplied using sterile stock solutions included S^0^, Na_2_S_2_O_3_·5H_2_O, Na_2_SO_3_, sulfate (MgSO_4_·7H_2_O), nitrate (NaNO_3_), ferric ion (Fe-citrate·4H_2_O), arsenate (Na_2_HAsO_4_·7H_2_O), NaAsO_2_, selenate (Na_2_SeO_4_), and Na_2_SeO_3_. Alternative terminal electron acceptor experiments were performed under anoxic conditions, with resazurin (2 mg/L) used as a visual indicator of anaerobic conditions and ascorbic acid (0.02%, w/v) as a reducing agent. Limited growth was observed using S^0^, Na_2_S_2_O_3_·5H_2_O, Na_2_SO_3_, NaNO_3_, Na_2_HAsO_4_·7H_2_O, and Na_2_SeO_4_ as electron acceptors (Table 1), while optimal growth occurred with O_2_ as the terminal electron acceptor across a range of concentrations. Oxygen sensitivity was determined from 0 to 21% (v/v) by injecting defined volumes of 100% (v/v) O_2_ into the headspace. Hydrogenotrophic growth occurred across the entire range, with optimal growth observed at 21% (v/v) O_2_ (Table 1). Neither S^0^ nor Na_2_S_2_O_3_·5H_2_O were required for growth.

Prior to these experiments, we experimentally determined KUI-RB^T^ could use neither ammonium (NH_4_Cl) nor sulfate (MgSO_4_·7H_2_O) as a sole electron donor or electron acceptor, respectively. Consequentially, these two compounds were retained in the basal medium across all experiments to supply nitrogen and sulfur.

### Morphology and chemotaxonomy

Cells were regularly viewed using phase contrast imaging on a Zeiss Primo Star light microscope at a magnification of 1000×. KUI-RB^T^ cells are motile, straight rod-shaped, and often found singularly or in pairs end-to-end. A Gram stain performed following standard protocols (MacFaddin, 2000) indicated that strain KUI-RB^T^ is Gram-negative.

Transmission electron microscopy was performed by AgResearch (Lincoln, New Zealand) on negatively stained and thin section cells of strain KUI-RB^T^ (Fig. 1). Negative stain images show KUI-RB^T^ to have peritrichous flagella (Fig. 1). Thin section images confirmed the presence of a plasma membrane, periplasmic space, and outer membrane typical of diderm bacteria (Fig. S2). No internal stacked membranes were observed in thin section images. Cells were measured as approximately 0.7 µm wide by 1.0 to 1.5 µm long.

**Fig. 1.**
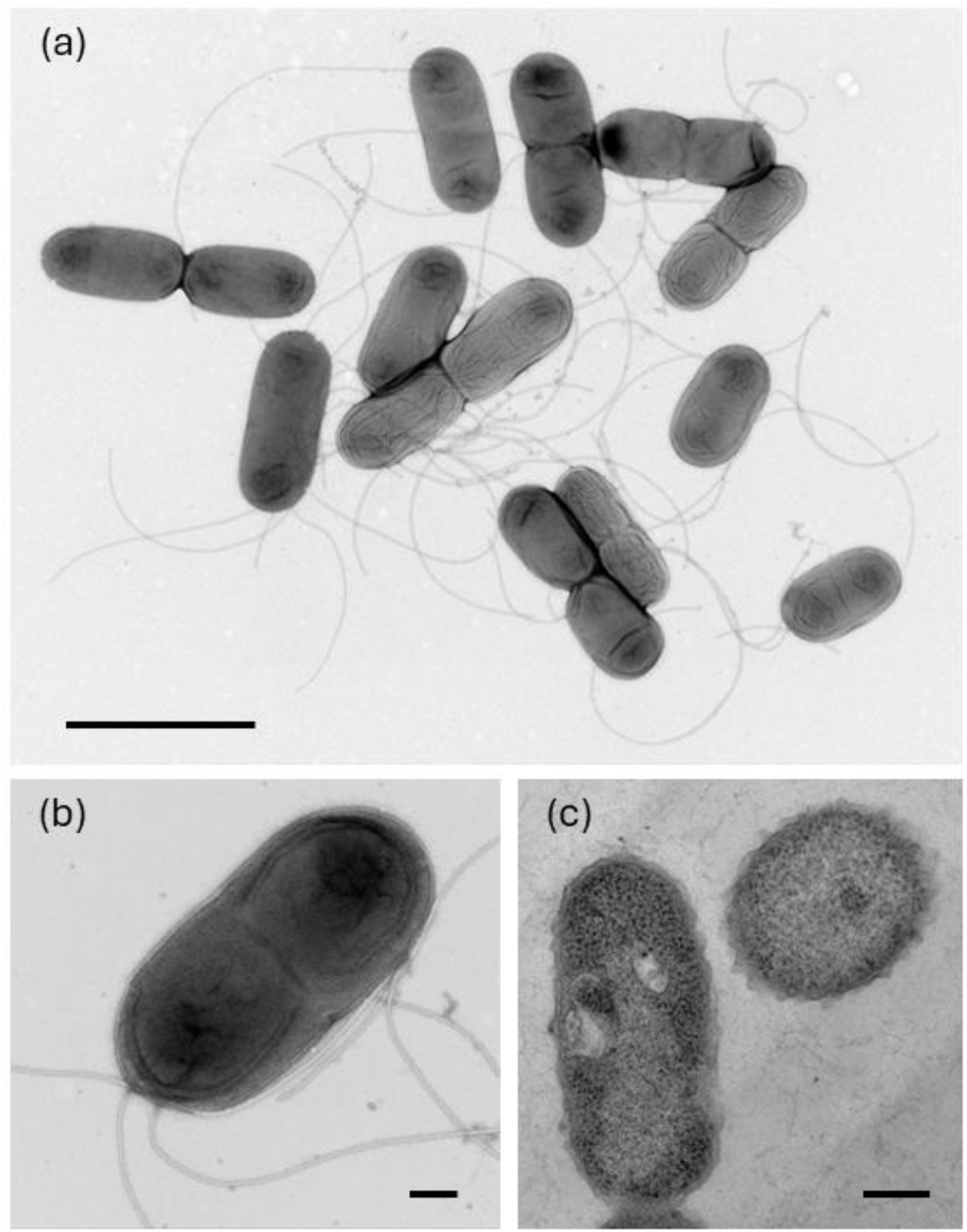
Transmission electron microscopy images of negatively stained (a, b) and thin section (c) cells of strain KUI-RB^T^. Scale bars represent 2 µm (a) and 200 nm (b, c).

Fatty acid and quinone analyses were performed by Callaghan Innovation (Wellington, New Zealand). Fatty acid methyl esters were prepared without lipid extraction as described by Svetashev *et al*. (1995). Fatty acids were analysed using an Agilent 7890 gas chromatograph with a flame ionisation detector and polar column (DB-wax UI, 30 m, 0.32 mm i.d.). The main fatty acids in KUI-RB^T^ are C_20:1_, C_18:1_, and C_18:0_ (Table S1). Respiratory quinones were extracted and analysed using a modified version of methodology described by Nishijima *et al*. (1997). The primary quinone in KUI-RB^T^ is MTK-7.

Metabolic characterisation experiments performed following standard protocols (MacFaddin, 2000) revealed KUI-RB^T^ to be catalase negative, oxidase positive, indole negative, and nitrate reductase positive. An API ZYM general enzyme activity kit (bioMérieux, Marcy-l’Étoile, France) indicated activity of alkaline phosphatase, C4 esterase, C8 esterase, leucine arylamidase, valine arylamidase, acid phosphatase, and napthol-AS-BI-phosphohydrolase in strain KUI-RB^T^.

### Genomics and phylogeny

Genomic DNA was extracted from cell pellets using the DNeasy PowerSoil Pro Kit (QIAGEN, Hilden, Germany). The quick-start protocol was followed as described, with elution performed by two sequential additions of 30 µL of PCR-grade water. Extracted DNA was cleaned and concentrated using a DNA Clean and Concentrator 25 Kit (Zymo Research, Irvine, USA). The kit protocol was followed as written, except spin columns IC were used in place of spin columns IICR and elution was performed by two sequential additions of 15 µL of PCR-grade water preheated to 70 °C. DNA concentration was determined using a Qubit dsDNA HS Assay Kit (Thermo Fisher Scientific, Waltham, USA) and DNA purity was assessed via NanoDrop. The DNA was stored at -20 °C and transported on ice to SeqCenter (Pittsburgh, USA) for whole genome sequencing. Illumina sequencing libraries were prepared using a tagmentation-based and PCR-based Illumina DNA Prep kit and custom IDT 10 bp unique dual indices with a target insert size of 280 bp. Illumina sequencing was performed on an Illumina NovaSeq X Plus sequencer producing 2×151 bp paired-end reads. Demultiplexing, quality control and adapter trimming were performed with bcl-convert1 (v4.2.4). Processing of whole genome sequencing data was performed in KBase (Arkin *et al*., 2018). Paired-end library reads were trimmed with Trimmomatic (v0.36) and assembled with Unicycler (v0.4.8), resulting in 37 contigs. The completeness and contamination of the genome is 99.19% and 0.20%, respectively, as determined by CheckM (v1.0.18). The whole genome is 1621227 bp with a G+C content of 34.23 mol%. The assembled contigs were uploaded to IMG and the genome was annotated with the IMG Annotation Pipeline (v.5.3.0) (IMG genome ID: 8136898124). The assembled contigs were also uploaded to the National Center for Biotechnology Information (NCBI; https://www.ncbi.nlm.nih.gov/) and annotated with PGAP (JBVODP000000000). The full-length 16S rRNA gene was extracted from the IMG annotated whole genome and uploaded to NCBI (PZ052650).

The full-length 16S rRNA gene sequence extracted from the whole genome was queried in BLAST using the core nucleotide database and default parameters (Altschul *et al*., 1990). A maximum-likelihood phylogenetic tree was generated with 16S rRNA gene sequences accessed through NCBI from all characterised members of the family *Hydrogenothermaceae* and a selection of closely related clones to identify the relationship of strain KUI-RB^T^ to other members of the phylum *Aquificota*. Sequences were aligned in ARB, then the phylogenetic tree was built with TREE-PUZZLE with the maximum-likelihood quartet-puzzling methodology, 10,000 puzzling steps, and ARB default parameters (Ludwig *et al*., 2004; Schmidt & von Haeseler, 2007). The phylogenetic placement of strain KUI-RB^T^ shows that it groups separately from representative *Sulfurihydrogenibium* and *Venenivibrio* with high confidence (quartet puzzling support value 95%; Fig. 2).

**Fig. 2.**
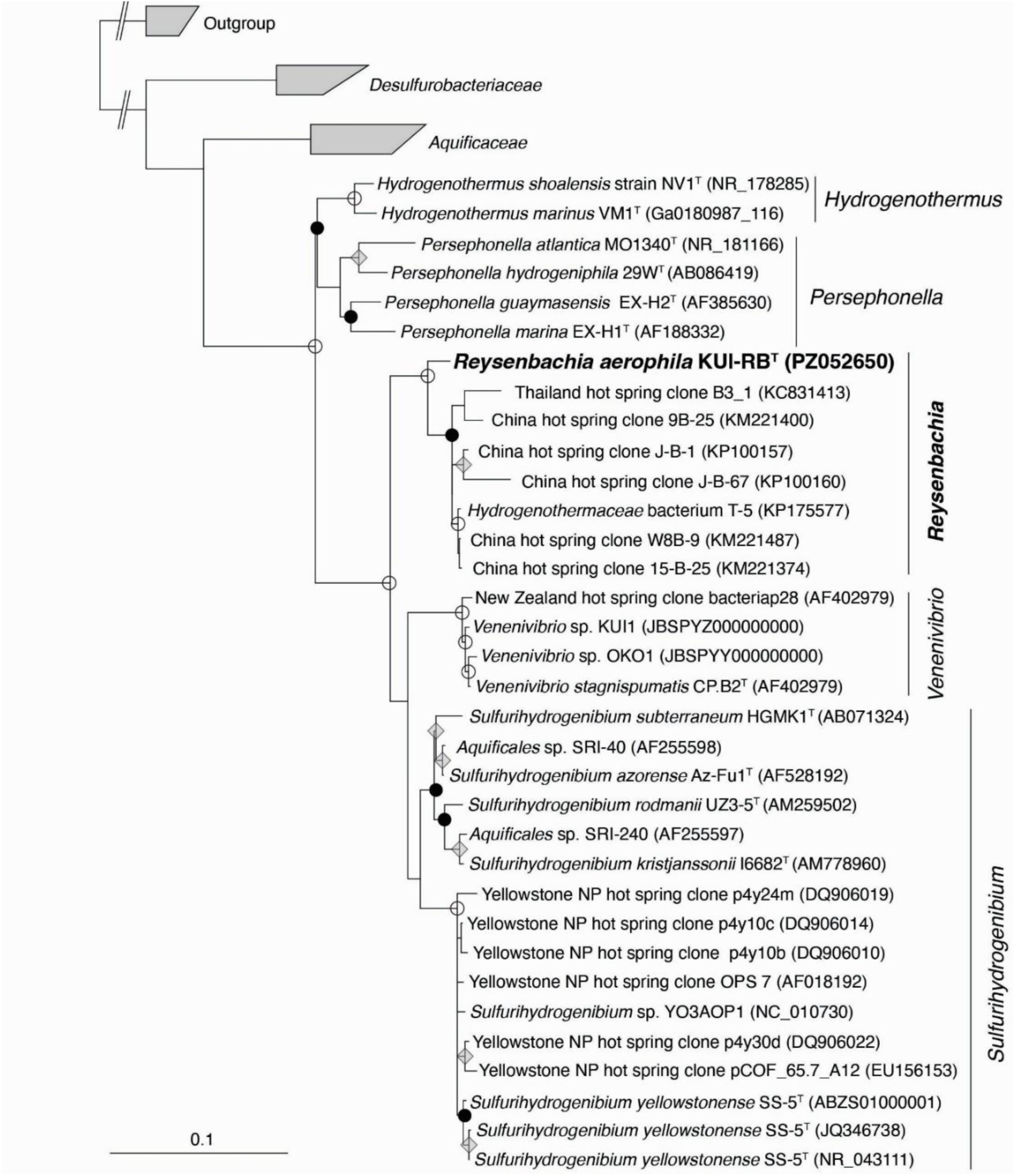
Maximum-likelihood phylogenetic tree based on 16S rRNA gene sequences, identifying the relationship between strain KUI-RB^T^ and other members of the phylum *Aquificota*. Sequences were aligned in ARB. The tree was built by TREE-PUZZLE with the maximum-likelihood quartet-puzzling methodology, 10,000 puzzling steps, and ARB default parameters. NCBI accession numbers are given in parentheses. The outgroup used to root the tree was composed of *Methanocaldococcus indicus* SL43^T^ (AF547621), *Methanocaldococcus jannaschii* JCM10045 (AB603516), and *Methanocaldococcus lauensis* SG7^T^ (MK602249) (not shown). *Venenivibrio* sp. KUI1 and *Venenivibrio* sp. OKO1 are unpublished isolates. Closed circles represent confidence >90%, open circles represent confidence >80%, grey diamonds represent confidence >70%. Bar represents 10 substitutions per 100 nucleotides.

To confirm these results, a maximum-likelihood phylogenomic tree was created using all available whole genome sequences of type strains in the family *Hydrogenothermaceae*, alongside the genome of strain KUI-RB^T^ and a *Hydrogenothermaceae* MAG (GCA_037478615, Slobodkina *et al*., 2025) that appears closely related to KUI-RB^T^. The phylogenomic tree was built using the Build Microbial SpeciesTree (v1.6.0) app in KBase (Arkin *et al*., 2018). This app utilized a set of 49 core universal genes, defined by Clusters of Orthologous Groups gene families, to perform multiple sequence alignment across the genomes. Alignments were then trimmed using GBLOCKS and the tree was built by FastTree2 (v2.1.10) with the fastest setting.

As with the 16S rRNA phylogenetic tree, the phylogenomic tree provided strong support for KUI-RB^T^ as the type strain of a novel species and genus, with a monophylogenetic clustering of KUI-RB^T^ and Uzbekistan MAG (GCA_037478615) (Fig. S3). Furthermore, these whole genomes were compared using FastANI (v0.1.3) and FastAAI (v0.1.20) and the nearest characterised species to strain KUI-RB^T^ had an average nucleotide identity (ANI) of 78.01% (Table S2) and an average amino acid identity (AAI) of 61.35% (Table S3), highlighting that strain KUI-RB^T^ is genomically distinct from all characterised *Hydrogenothermaceae* (Gerhardt *et al*., 2025; Jain *et al*., 2018).

### Comparison with related genera

The relatively small genome and low G+C content of strain KUI-RB^T^ is typical of the family *Hydrogenothermaceae* (Table S4). Phylogenetic analyses of the 16S rRNA gene demonstrate that KUI-RB^T^ is a member of the family *Hydrogenothermaceae*, with *S. azorense* Az-Fu1^T^ as its closest characterised relative (94.51% similarity). The closest cultured relative of strain KUI-RB^T^ is *Hydrogenothermaceae* strain T-5, isolated from a white streamer community in a sulfurous seep (70 °C, pH 6.6) in Tengchong, China (Hedlund *et al*., 2015). BLAST pairwise analysis (Altschul *et al*., 1990) of the KUI-RB^T^ 16S rRNA gene sequence identified eight clone sequences with sequence similarities ranging from 95-97%, originating from geothermal springs in Tengchong, China and Betong, Thailand (Hedlund *et al*., 2015; Li *et al*., 2016). These sequences form a monophyletic group, suggesting this cluster of strains is globally distributed in terrestrial geothermal springs (Fig. 2).

A survey of MAGs and genomes found a closely related MAG (GCA_037478615) assembled from a microbial community sampled from a hot stream sediment in the Navoiy region of Uzbekistan (Slobodkina *et al*., 2025). ANI and AAI analyses of these two genomes (99.35% and >90%, respectively; Table S2, Table S3) suggest that GCA_037478615 and KUI-RB^T^ are strains of the same species. ANI of strain KUI-RB^T^ against all other *Hydrogenothermaceae* genomes ranged from 76 to 78% (Table S2) and AAI was 53 to 61% (Table S3), further supporting strain KUI-RB^T^ as a novel species in a new genus of *Hydrogenothermaceae*.

The temperature, pH and salinity growth ranges and optima of strain KUI-RB^T^ are comparable to those of other members of the family *Hydrogenothermaceae* (Table S4). However, KUI-RB^T^ is the only member of the family observed to grow optimally under atmospheric oxygen conditions (21% v/v); all other cultured *Hydrogenothermaceae* are microaerophilic, with reported optimal growth at < 15% (v/v) O_2_ (Table S4). The genome of KUI-RB^T^ contains genes encoding cytochrome b (ACZ6J7_05150) and cytochrome c (ACZ6J7_05255), as is observed in all other *Hydrogenothermaceae* genomes (GCF_000021545, GCF_000021565, GCF_000173615, GCF_016617615, GCF_003688665, GCF_000619805, GCF_900215515, GCF_900182795).

KUI-RB^T^ was able to utilize a wide range of organic carbon substrates as carbon sources in the presence of H_2_, but could not use them as energy sources, similar to *S. azorense* and *S. yellowstonense* (Table 1). H_2_ was the only electron donor used by KUI-RB^T^, as seen in *V. stagnispumantis* and *H. marinus* (Table S4). This use of H_2_ as an energy source is mediated by Ni/Fe hydrogenases encoded in the genome of strain KUI-RB^T^ (ACZ6J7_08490, ACZ6J7_08590, ACZ6J7_04585). The use of ammonium as a source of nitrogen is also supported, with multiple ammonium transporters in the genome (ACZ6J7_05590, ACZ6J7_07840).

KUI-RB^T^ is the only known member of the family to utilize thiosulfate and sulfite as sole electron acceptors, and, like *S. azorense* and *P. marina*, it can utilize elemental sulfur as a sole electron acceptor (Table S4) (Aguiar *et al*., 2004; Götz *et al*., 2002). While there is a YeeE/YedE thiosulfate transporter family protein (ACZ6J7_04805) in the genome of KUI-RB^T^, there are no annotated thiosulfate reductases, therefore it is hypothesized that thiosulfate disproportionates to elemental sulfur and sulfite in the system, which are then used as terminal electron acceptors. The genome of KUI-RB^T^ encodes sulfur oxidation genes *soxA* (ACZ6J7_04900), *soxB* (ACZ6J7_04895), *soxX* (ACZ6J7_04920), *soxY* (ACZ6J7_04910), and *soxZ* (ACZ6J7_04905), but unlike *Sulfurihydrogenibium* and *Persephonella* spp., growth of strain KUI-RB^T^ was not observed using reduced sulfur compounds as electron donors in the absence of H_2_ (Table S4). The presence of thiosulfate or elemental sulfur was not required for growth of KUI-RB^T^ as it is for *V. stagnispumantis* (Hetzer *et al*., 2008), and we did not observe enhanced growth of KUI-RB^T^ when thiosulfate or elemental sulfur was present with H_2_, as was reported in *S. azorense* (Aguiar *et al*., 2004). This contradiction between the presence of sulfur oxidation genes and apparent lack of sulfur oxidation is a potential line of investigation for future research.

Arsenate, selenate, and nitrate were also used by KUI-RB^T^ as alternative electron acceptors, which has been observed in other *Hydrogenothermaceae* (Table S4) (Aguiar *et al*., 2004; François *et al*., 2021; Götz *et al*., 2002; Nakagawa *et al*., 2003; Takai *et al*., 2003). Supporting these phenotypic observations, the genome of KUI-RB^T^ encodes an arsenate reductase ArsC (ACZ6J7_02235), although no nitrate-reductase or selenate-reductase genes were identified in the genome. KUI-RB^T^ did not utilize arsenite as an electron donor or acceptor (Table 1), which aligns with the lack of annotated arsenite oxidase gene in its genome. However, mechanisms of arsenic detoxification are evident in the genome of KUI-RB^T^ through an arsenic transporter (ACZ6J7_02240) and two ArsR/SmtB family transcription factors (ACZ6J7_02220, ACZ6J7_02405). Detoxification of arsenic has been reported previously in *V. stagnispumantis*, as evidenced by its growth in elevated arsenite and arsenate levels (Hetzer, 2007).

KUI-RB^T^ cells are motile, Gram-negative rods, similar in size to other *Hydrogenothermaceae*. KUI-RB^T^ have peritrichous flagella like *S. azorense* (Aguiar *et al*., 2004), while most other *Hydrogenothermaceae* have polar flagella (François *et al*., 2021; Nakagawa *et al*., 2005; O’Neill *et al*., 2008; Stohr *et al*., 2001; Takai *et al*., 2003). No sporulation of KUI-RB^T^ was observed; however, when cells were no longer viable, they appeared as translucent cocci approximately 1.0 to 1.5 µm in diameter with a denser inner circle, as reported previously in *P. atlantica* (François *et al*., 2021). No internal stacked membranes were observed in thin section imaging, as has been reported in *S. azorense*, *S. kristjanssonii*, and *P. marina* (Aguiar *et al*., 2004; Flores *et al*., 2008; Götz *et al*., 2002). The primary quinone of KUI-RB^T^ is MTK-7, which is consistent with other *Hydrogenothermaceae*. With respect to fatty acid content, KUI-RB^T^ possessed a fatty acid suite that generally reflected those of other *Hydrogenothermaceae*; although, it included a greater proportion of C_18:1_ than C_18:0_, a unique feature amongst *Hydrogenothermaceae* (Table S1).

In conclusion, we isolated a facultatively anaerobic, hydrogen-oxidizing, thermophilic bacterium from biofilm collected from a geothermal spring in Kuirau Park, Rotorua, New Zealand. This bacterium, strain KUI-RB^T^, utilizes hydrogen as an electron donor and carbon dioxide and various organic carbon substrates as carbon sources. KUI-RB^T^ grows optimally using oxygen as a terminal electron acceptor, but can utilize elemental sulfur, thiosulfate, sulfite, nitrate, arsenate, and selenate as alternative terminal electron acceptors. Based on the phylogenetic, phylogenomic, and physiological characterisations presented here, we propose KUI-RB^T^ as a novel species in a new genus of *Hydrogenothermaceae*, for which we propose the name *Reysenbachia aerophila* gen. nov., sp. nov. The type strain is KUI-RB^T^ (=KCTC accession =JCM accession).

### Description of *Reysenbachia* gen. nov

*Reysenbachia* [Rey.sen.ba’chi.a N.L. fem n. named in honour of the microbiologist Anna-Louise Reysenbach, for her substantial contributions to the study of *Aquificales* and for pushing boundaries as a pioneering woman in extremophile microbiology].

Motile Gram-negative rods. Thermophilic and neutrophilic. Facultatively anaerobic and facultatively heterotrophic. Capable of chemolithoautotrophic and chemolithoheterotrophic growth. NaCl is not required for growth. Major fatty acids include C_20:1_, C_18:1_, and C_18:0_. Primary quinone is MTK-7. Most closely related to genera *Sulfurihydrogenibium* and *Venenivibrio* based on 16S rRNA gene sequence and whole genome analyses. Occurs in terrestrial geothermal features. Type species is *Reysenbachia aerophila*.

### Description of *Reysenbachia aerophila* sp. nov

*Reysenbachia aerophila* [ae.ro’phi.la. Gr. n. *aēr*, air; N.L. fem. adj. *aerophila*, air-loving].

Exhibits the following properties, in addition to those described for the genus. Cells are straight, motile rods, measuring approximately 0.7 µm by 1.0 to 1.5 µm with peritrichous flagella.

Growth was observed from 39 °C to 74 °C, pH 5.0 to pH 7.5, 0 to 1% (w/v) NaCl, and 0 to 21% (v/v) oxygen, and optimum growth was observed at 64.5 °C, pH 6.5, 0.4-0.7% (w/v) NaCl, and 21% (v/v) oxygen, respectively. Can utilize hydrogen as an electron donor. Can utilize oxygen, elemental sulfur, thiosulfate, sulfite, nitrate, arsenate, and selenate as electron acceptors. Can utilize carbon dioxide, Bacto peptone, trypticase peptone, Casamino acids, sucrose, glucose, starch, formate, citrate, succinate, acetate, and pyruvate as carbon sources, but not energy sources. Catalase negative, oxidase positive, indole negative, nitrate reductase positive. Notable activity of alkaline phosphatase, C4 esterase, C8 esterase, leucine arylamidase, valine arylamidase, acid phosphatase, and napthol-AS-BI-phosphohydrolase. The 16S rRNA gene sequence is 94.51% similar to that of *S. azorense* Az-Fu1^T^. The whole genome has an ANI of 78.01% to *S. azorense* Az-Fu1^T^ and an AAI of 61.35% to *S. subterraneum* HGMK1^T^.

The type strain, KUI-RB^T^ (=KCTC accession =JCM accession), was isolated from biofilm, collected from a geothermal spring in Kuirau Park, Rotorua, New Zealand. The DNA G+C content for the type strain is 34.23 mol%.

## Acknowledgements

The authors would like to thank Professor Anna-Louise Reysenbach for her guidance and mentorship. The authors also wish to thank Marina Richena, Kim Parker, and Duane Harland for their assistance with transmission electron microscopy performed at AgResearch. Thank you to Rotorua District Council for permission to sample at Kuirau Park. This work was supported by a New Zealand Royal Society Marsden grant (MFP-UOC2201).

## Supplementary figures and tables

**Fig. S1.**
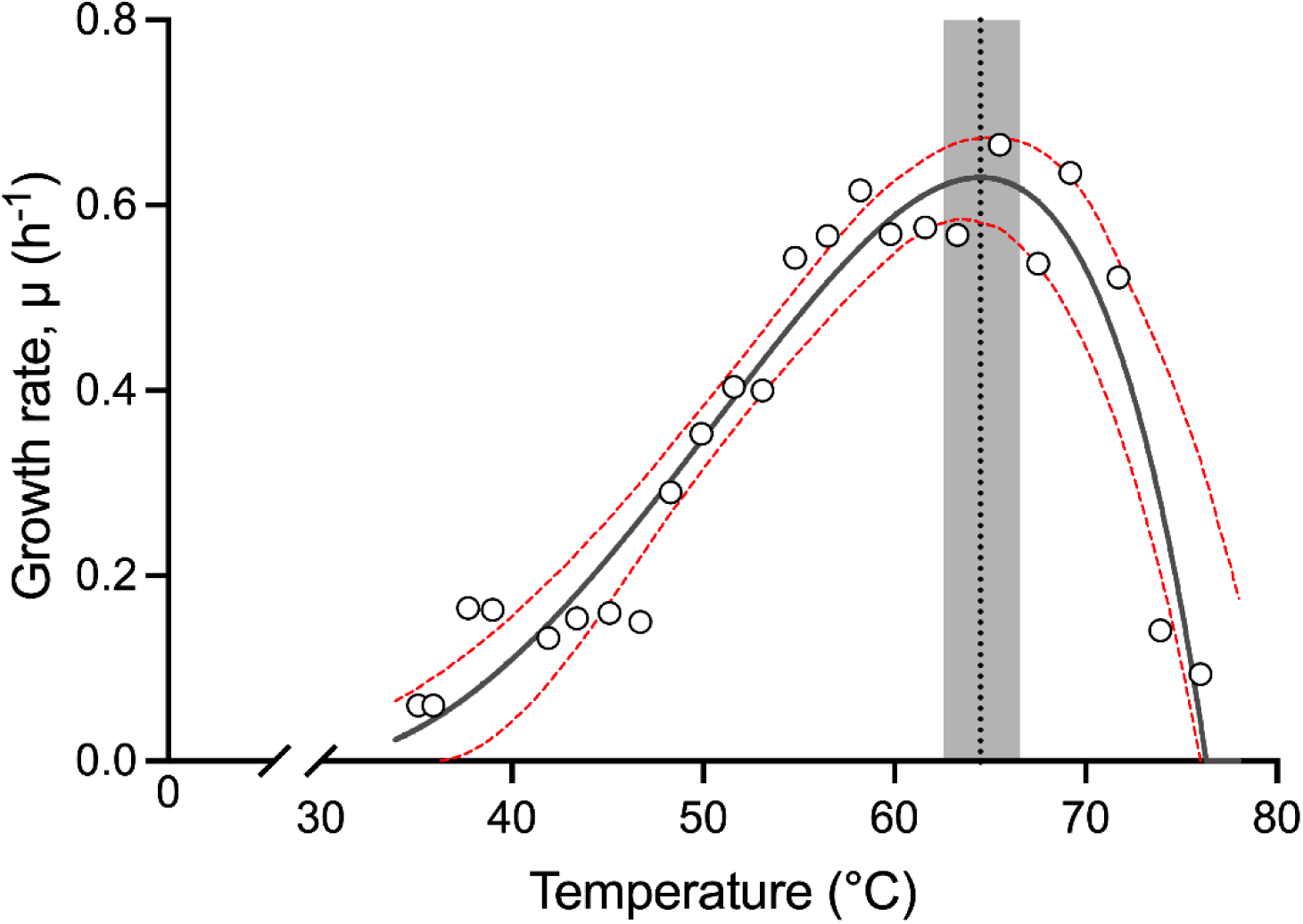
Temperature dependence of specific growth rate (μ, h^-1^) of strain KUI-RB^T^ modelled using the Rosso cardinal temperature model with inflection. Open circles represent observed growth rates. The fitted model is shown as a solid black line, with red dashed lines indicating the 95% confidence interval of μ across the temperature range. Predicted optimum temperature (T_opt_) is indicated by a vertical dashed line, and shaded region represents the 95% confidence interval of T_opt_ derived from 2,000 bootstrap replicates.

**Fig. S2.**
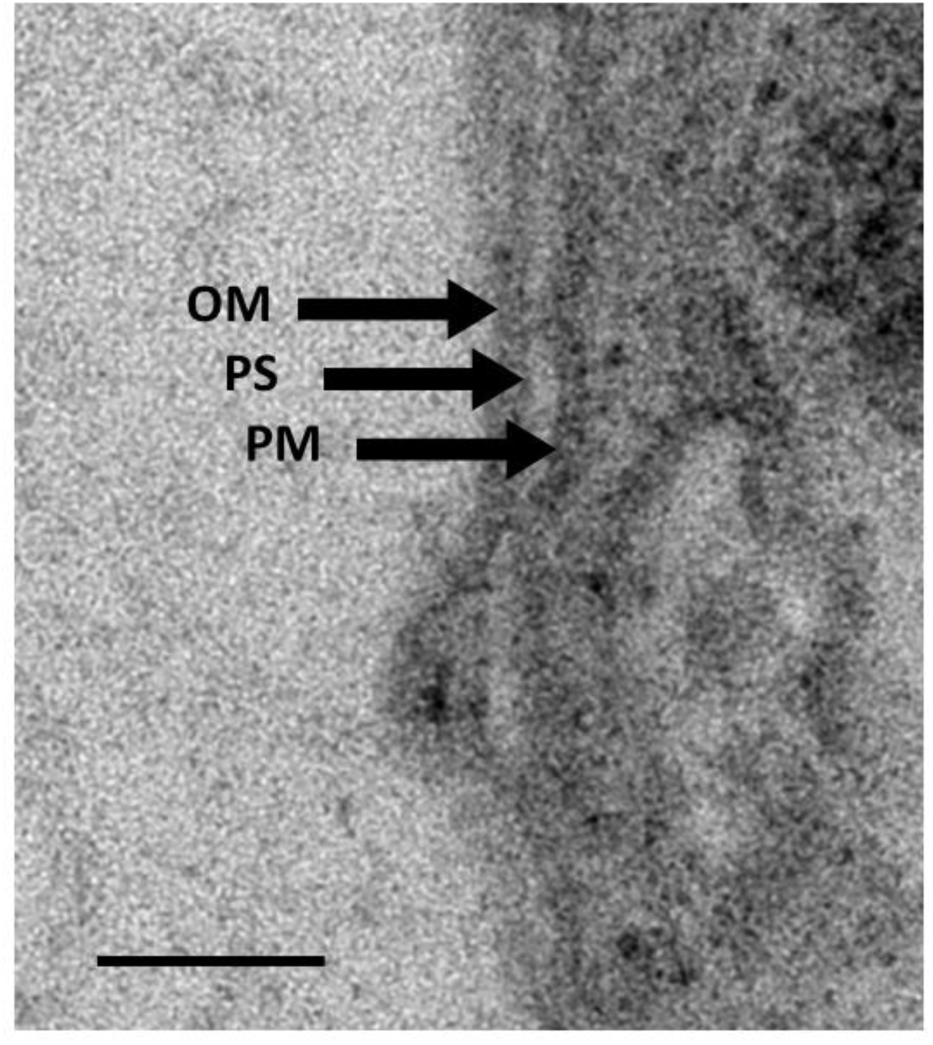
Transmission electron microscopy of thin section cell membrane of strain KUI-RB^T^. OM, outer membrane; PS, periplasmic space; PM, plasma membrane. Scale bar represents 50 nm.

**Fig. S3.**
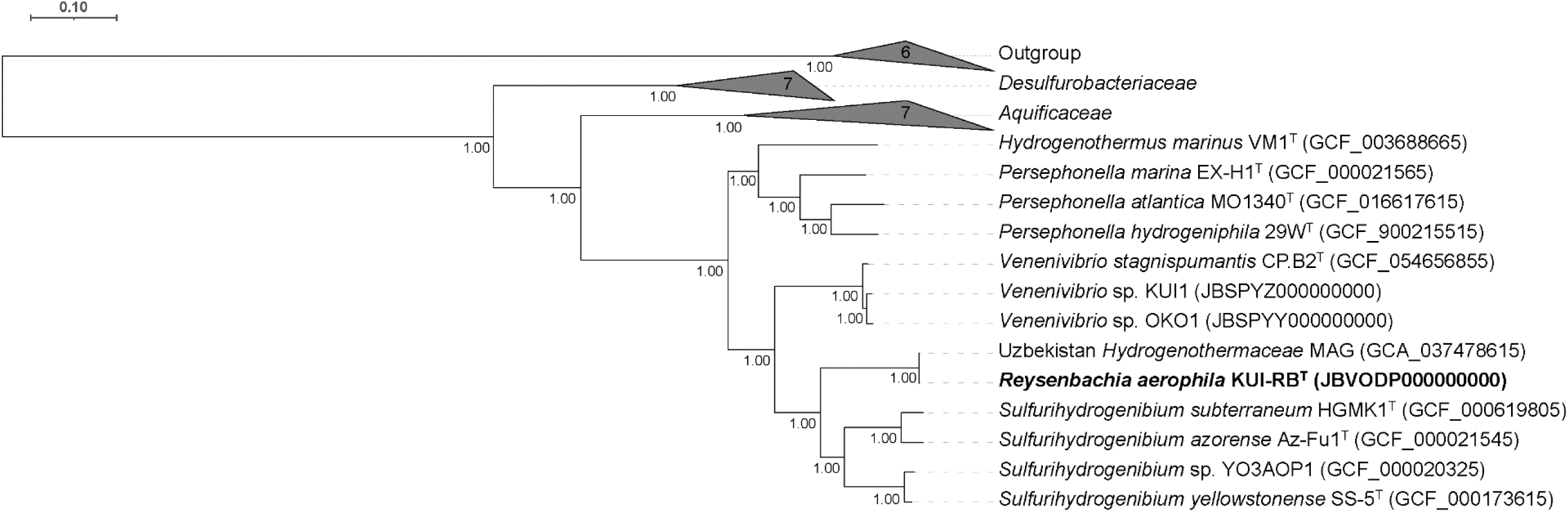
Maximum-likelihood phylogenomic tree based on whole genome sequences, identifying the relationship between strain KUI-RB^T^ and other members of the phylum *Aquificota*. In the Build Microbial SpeciesTree (v1.6.0) app in KBase, a set of 49 core universal genes was used to perform multiple sequence alignment across the genome and alignments were trimmed using GBLOCKS. The tree was then built by FastTree2 (v2.1.10) with the fastest setting. NCBI accession numbers are given in parentheses. The outgroup used to root the tree was composed of *Methanocaldococcus bathoardescens* JH146^T^ (GCF_000739065.1), *Methanocaldococcus fervens* AG86^T^ (GCF_000023985.1), *Methanocaldococcus infernus* ME^T^ (GCF_000092305.1), *Methanocaldococcus jannaschii* DSM 2661^T^ (GCF_000091665.1), *Methanocaldococcus villosus* KIN24-T80^T^ (GCF_000371805.1), and *Methanocaldococcus vulcanius* M7^T^ (GCF_000024625.1) (not shown). *Venenivibrio* sp. KUI1 and *Venenivibrio* sp. OKO1 are unpublished isolates. Bar represents 10 substitutions per 100 nucleotides.

**Table S1.** Comparison of fatty acid composition of strain KUI-RB^T^ with type strains of the family *Hydrogenothermaceae*. Species: 1, *Reysenbachia aerophila* KUI-RB^T^ (this study); 2, *Venenivibrio stagnispumantis* CP.B2^T^ (this study); 3, *Sulfurihydrogenibium subterraneum* HGMK1^T^ (Takai *et al*., 2003); 4, *Hydrogenothermus marinus* (VM1^T^) (Stohr *et al*., 2001); 5, *Persephonella guaymasensis* (EX-H2^T^) (Götz *et al*., 2002); 6, *Persephonella hydrogeniphila* (29W^T^) (Nakagawa *et al*., 2003); 7, *Persephonella marina* (EX-H1^T^) (Götz *et al*., 2002).

| Fatty acid | 1 | 2 | 3 | 4 | 5 | 6 | 7 |
| --- | --- | --- | --- | --- | --- | --- | --- |
| C<sub>12:0</sub> | 1.3 | 2.9 | 3.1 |  |  |  |  |
| C<sub>14:0</sub> |  |  |  |  |  |  |  |
| C<sub>16:0</sub> | 4.0 | 1.6 | 1.9 |  |  | 4.2 |  |
| C<sub>16:1</sub> | 1.1 |  |  |  |  |  |  |
| C<sub>18:0</sub> | 10.7 | 31.1 | 25.4 | 23.0 | 26.3 | 24.7 | 16.3 |
| C<sub>18:1</sub> | 33.3 | 12.0 | 13.6 | 15.5 | 12.6 | 23.8 | 16.0 |
| C<sub>20:0</sub> | 0.5 | 4.2 | 2.3 |  | 1.8 |  | 6.1 |
| C<sub>20:1</sub> | 49.1 | 48.0 | 53.7 | 48.0 | 45.0 | 40.8 | 21.0 |
| Total | 100.0 | 99.8 | 100.0 | 86.5 | 85.7 | 93.5 | 59.4 |

**Table S2.** Whole genome average nucleotide identity of strain KUI-RB^T^ in comparison to type strains of the family *Hydrogenothermaceae*. Genomes are listed in rank order. ANI analysis was undertaken using FastANI (v0.1.3). NCBI accession numbers are given in parentheses.

| Query | Reference | ANI (%) |
| --- | --- | --- |
| *Reysenbachia aerophila*<br>KUI-RB<sup>T</sup><br>(JBVODP000000000) | Uzbekistan hot spring MAG (GCA_037478615) | 99.3459 |
|  | *Sulfurihydrogenibium azorense* Az-Fu1<sup>T</sup> (GCF_000021545) | 78.0107 |
|  | *Sulfurihydrogenibium subterraneum* HGMK1<sup>T</sup> (GCF_000619805) | 77.4155 |
|  | *Sulfurihydrogenibium yellowstonense* SS-5<sup>T</sup> (GCF_000173615) | 77.1609 |
|  | *Venenivibrio* sp. OKO1* (JBSPYY000000000) | 77.0617 |
|  | *Sulfurihydrogenibium* sp. YO3AOP1 (GCF_000020325) | 77.0445 |
|  | *Venenivibrio stagnispumantis* CP.B2<sup>T</sup> (GCF_054656855) | 76.9736 |
|  | *Venenivibrio* sp. KUI1* (JBSPYZ000000000) | 76.9227 |
|  | *Persephonella marina* EX-H1<sup>T</sup> (GCF_000021565) | 76.5020 |
|  | *Hydrogenothermus marinus* VM1<sup>T</sup> (GCF_003688665) | 76.3872 |
|  | *Persephonella atlantica* MO1340<sup>T</sup> (GCF_016617615) | 76.1333 |
|  | *Persephonella hydrogeniphila* 29W<sup>T</sup> (GCF_900215515) | 75.9698 |

\* *Venenivibrio* sp. OKO1 and *Venenivibrio* sp. KUI1 are unpublished isolates.

**Table S3.** Whole genome average amino acid identity of strain KUI-RB^T^ in comparison to type strains of the family *Hydrogenothermaceae*. Single copy proteins (SCPs) within the genomes were compared using FastAAI (v0.1.20). NCBI accession numbers are given in parentheses.

| Query | Target | AAI (%) | Average Jaccard Index | Shared SCPs | Maximum SCPs |
| --- | --- | --- | --- | --- | --- |
| *Reysenbachia aerophila* KUI-RB<sup>T</sup> (JBVODP000000000) | Uzbekistan hot spring MAG (GCA_037478615) | >90 | 0.9725 | 76 | 76 |
|  | *Sulfurihydrogenibium subterraneum* HGMK1<sup>T</sup> (GCF_000619805) | 61.35 | 0.3704 | 75 | 75 |
|  | *Sulfurihydrogenibium azorense* Az-Fu1<sup>T</sup> (GCF_000021545) | 61.34 | 0.3703 | 74 | 74 |
|  | *Sulfurihydrogenibium* sp. YO3AOP1 (GCF_000020325) | 60.53 | 0.3539 | 75 | 75 |
|  | *Venenivibrio* sp. KUI1* (JBSPYZ000000000) | 58.69 | 0.3171 | 74 | 74 |
|  | *Venenivibrio stagnispumantis* CP.B2<sup>T</sup> (GCF_054656855) | 58.66 | 0.3165 | 74 | 74 |
|  | *Sulfurihydrogenibium yellowstonense* SS-5<sup>T</sup> (GCF_000173615) | 58.61 | 0.3154 | 56 | 56 |
|  | *Venenivibrio* sp. OKO1* (JBSPYY000000000) | 58.55 | 0.3143 | 75 | 75 |
|  | *Persephonella atlantica* MO1340<sup>T</sup> (GCF_016617615) | 54.07 | 0.2291 | 75 | 76 |
|  | *Persephonella hydrogeniphila* 29W<sup>T</sup> (GCF_900215515) | 54.07 | 0.2291 | 76 | 76 |
|  | *Persephonella marina* EX-H1<sup>T</sup> (GCF_000021565) | 54.02 | 0.2281 | 76 | 76 |
|  | *Hydrogenothermus marinus* VM1<sup>T</sup> (GCF_003688665) | 53.30 | 0.2153 | 72 | 74 |

\* *Venenivibrio* sp. KUI1 and *Venenivibrio* sp. OKO1 are unpublished isolates.

**Table S4.**
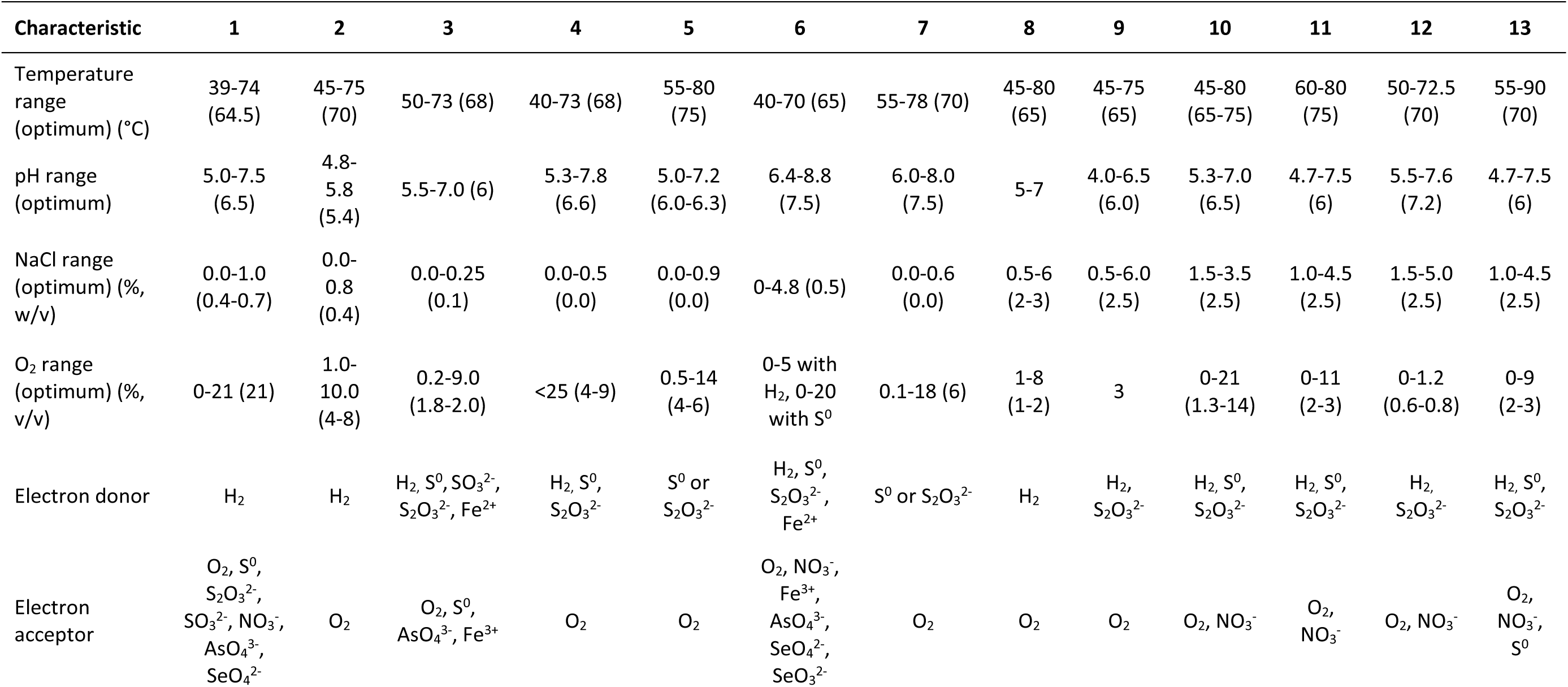
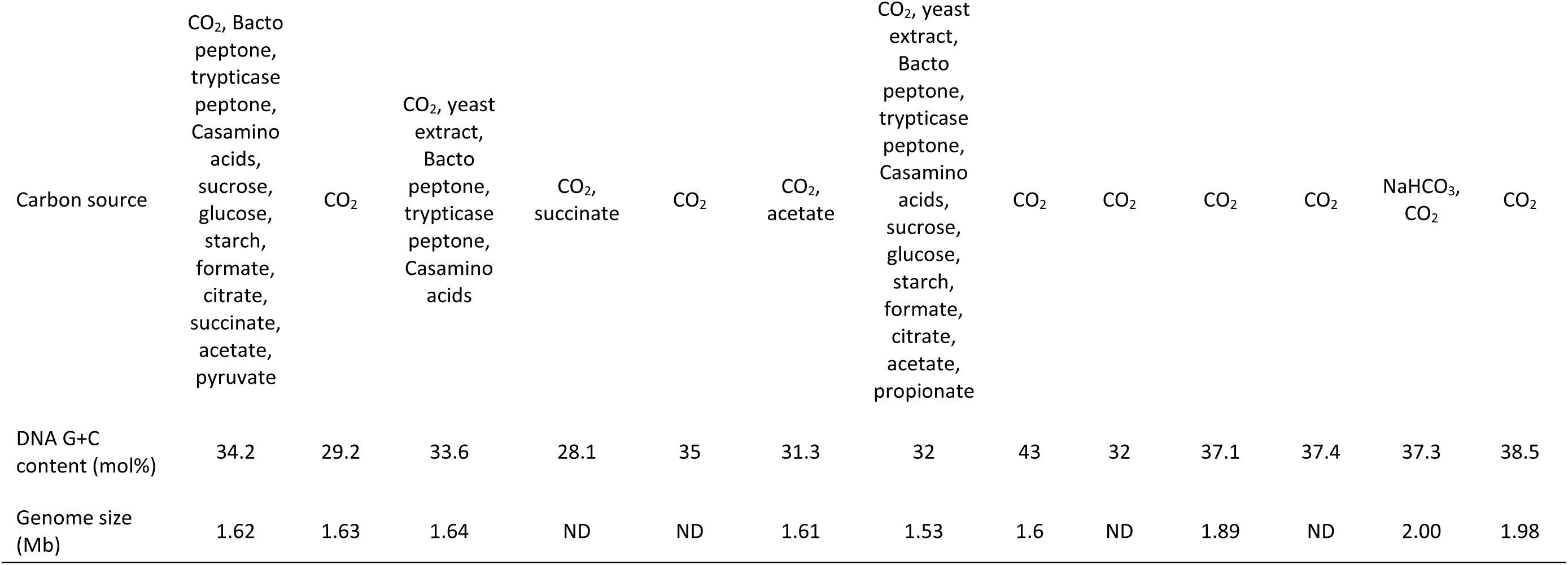
Differential characteristics of strain KUI-RB^T^ and expanded related type strains of the family *Hydrogenothermaceae*. Species: 1, *Reysenbachia aerophila* KUI-RB^T^ (this study); 2, *Venenivibrio stagnispumantis* CP.B2^T^ (Hetzer *et al*., 2008); 3, *Sulfurihydrogenibium azorense* AZ-Fu1^T^(Aguiar *et al*., 2004; Nakagawa *et al*., 2005); 4, *Sulfurihydrogenibium kristjanssonii* I6628^T^ (Flores *et al*., 2008); 5, *Sulfurihydrogenibium rodmanii* UZ3-5^T^(O’Neill *et al*., 2008); 6, *Sulfurihydrogenibium subterraneum* HGMK1^T^ (Aguiar *et al*., 2004; Nakagawa *et al*., 2005; Takai *et al*., 2002); 7, *Sulfurihydrogenibium yellowstonense* SS-5^T^ (Nakagawa *et al*., 2005); 8, *Hydrogenothermus marinus* (VM1^T^) (Stohr *et al*., 2001); 9, *Hydrogenothermus shoalensis* (NV1^T^) (Sislak, 2013); 10, *Persephonella atlantica* (MO1340^T^) (François *et al*., 2021); 11, *Persephonella guaymasensis* (EX-H2^T^) (Götz *et al*., 2002); 12, *Persephonella hydrogeniphila* (29W^T^) (Nakagawa *et al*., 2003); 13, *Persephonella marina* (EX-H1^T^) (Götz *et al*., 2002). ND, Not determined.

## Supplementary methods

### Transmission electron microscopy

Cells were negatively stained with 1% uranyl acetate on Formar 200-mesh copper grids (Proscitech, Thuringowa, Australia) that had been coated with a 3 nm carbon layer and charged by glow discharge treatment (Leica Mikrosysteme GmbH, Vienna, Austria). Cells were treated for thin sectioning using Karnovsky’s fixative (2% glutaraldehyde, 2% formaldehyde in 0.1 M sodium cacodylate buffer, pH 7.4) then rinsed repeatedly in 0.1 M sodium cacodylate buffer. Samples were resuspended in 3% ultra-low melting point agarose, cooled, and stored overnight in 0.1 M cacodylate buffer. Further fixation was performed in 1% osmium tetroxide in 0.1 M phosphate buffer (pH 7.4), followed by repeated rinsing in 0.1 M phosphate buffer then ultrapure water. Samples were dehydrated through a graded ethanol series then stored in acetone overnight. Samples were treated with increasing ratios of epoxy resin and 100% resin overnight. A final resin change was performed, then samples were cured at 60 °C. Ultrathin sections (90 nm) were cut using a Leica EM UC7 ultramicrotome and collected on Leica EM ACE600 glow-discharged formvar 200-mesh copper grids. Sample grids were stained with 2% uranyl acetate, dried, treated with 0.2% lead citrate in ultrapure water, and rinsed repeatedly in ultrapure water. Negative stain and thin section sample grids were analysed in a Talos 120C transmission electron microscope (Thermo Fisher Scientific, Waltham, USA). Images were taken using a Ceta camera (4096×4096) and the MAPS 3.27 software was used for montaging and large-scale mapping.

